# Lipid nanoparticle-encapsulated mRNA encoding CagW, a conserved component of the *Helicobacter pylori* type IV secretion system, protects mice against *H. pylori* challenge

**DOI:** 10.64898/2026.10.28.762502

**Authors:** Yunwang Zhao, Hongxian Cheng, Shiyi Zhou, Lei He, Chenlin Xiu, Baihai Huang, Xiongxiong Lu

**Author notes:** Correspondence: Xiongxiong Lu.

## Abstract

**Background:** Roughly one person in two carries *Helicobacter pylori*, a pathogen classified by the World Health Organization as a group I carcinogen, and the steady loss of antibiotic efficacy has made prophylaxis an urgent rather than an optional goal. No mRNA vaccine against this organism has yet been tested in the clinic, and CagW, a conserved inner-membrane subunit of the *H. pylori* type IV secretion system (T4SS), has never been evaluated as an mRNA-encoded antigen.

**Methods:** A codon-optimised construct carrying a signal peptide, the CagW coding sequence and a human IgG Fc tag was synthesised, transcribed into mRNA, encapsulated in lipid nanoparticles (LNPs) and shown to direct antigen expression in HEK293T cells. C57BL/6 mice received three intramuscular doses of CagW-Fc mRNA-LNPs (5 µg mRNA each) at 14-day intervals; animals given alum alone, luciferase mRNA-LNPs or CagW protein adsorbed to alum served as comparators. Humoral readouts (IgG, IgG1, IgG2a, IgM, IgA and secretory IgA), lymphocyte proliferation, CD4^+^/CD8^+^ subsets, intracellular IFN-γ, IL-4 and IL-17A and circulating cytokines were quantified before challenge, and protection was assessed after live challenge.

**Results:** The formulation (93.2 nm; PDI 0.13; approximately 90% encapsulation) produced CagW-specific protein in vitro. Vaccination raised high-titre CagW-specific IgG (maximal on day 42), IgM and mucosal sIgA, and drove Th1/Th2-type cellular activation (IL-2, IL-4, IL-12, IFN-γ and IL-17A). After challenge, immunised animals carried markedly fewer gastric bacteria and showed lower urease activity than controls.

**Conclusions:** CagW-Fc mRNA-LNPs are therefore a credible mRNA vaccine platform for *H. pylori* and warrant further preclinical development.

## Introduction

*Helicobacter pylori* is a microaerophilic, Gram-negative spiral bacterium that colonises the gastric mucus layer, and it remains the commonest chronic bacterial infection of humans: roughly one in two people worldwide is a carrier.^5,6^ Acquisition generally occurs in early childhood, and transmission is thought to proceed mainly along faecal–oral and oral–oral routes. Prevalence is strikingly uneven, with figures of 20– 40% in Western countries but 70–90% in parts of Asia and in many developing regions.^7^ Once established, colonisation persists for decades and sustains a chronic gastritis that may progress to atrophic gastritis, peptic ulceration, mucosa-associated lymphoid tissue (MALT) lymphoma and adenocarcinoma; *H. pylori*-driven inflammation is estimated to underlie about three-quarters of gastric cancers and 5.5% of all human malignancies,^8^ and infection-associated inflammatory burden tracks with excess all-cause mortality.^9^ These observations underpin the classification of the organism as a group I carcinogen.

Control of *H. pylori* still rests almost entirely on antibiotic-based eradication. Classical triple therapy, and the bismuth-containing quadruple regimens that have largely displaced it in many settings, pair two or three antimicrobials with a proton pump inhibitor (PPI) and, where used, a bismuth salt.^10^ Cure rates have fallen as resistance to clarithromycin, metronidazole and levofloxacin has spread, so that susceptibility-guided prescribing is now widely advocated.^11^ Recrudescence and genuine reinfection after apparently successful eradication further complicate long-term management and are particularly frequent in children.^12^ Practical obstacles compound the pharmacological ones: regimens are costly by the standards of the regions where infection is most prevalent, adverse effects erode adherence, and access to endoscopy-based or molecular diagnosis is limited, which has stimulated interest in low-cost biosensor platforms.^13^ Together these constraints explain why a prophylactic vaccine has long been regarded as the most rational route to reducing the global burden of *H. pylori*-related disease.

Vaccination has repeatedly proved the most cost-effective way of controlling mucosal pathogens, yet no licensed *H. pylori* vaccine exists. Recombinant subunit antigens, including UreB, heat-shock proteins, VacA, CagA and catalase, have reproducibly lowered bacterial loads in rodent models without achieving sterile protection, and monovalent protein formulations adsorbed to conventional adjuvants have proved particularly disappointing. DNA vaccines such as the cagW-based construct evaluated by Chehelgerdi and Doosti are immunogenic in BALB/c mice,^1,14^ but naked plasmid delivery is constrained by poor transfection efficiency and nuclease-mediated degradation, while immunoinformatics-led designs, including a recently reported self-amplifying RNA multi-epitope candidate, remain largely untested in vivo.^2^ Messenger RNA delivered in lipid nanoparticles (LNPs) offers an alternative: the ionisable lipid shell shields the transcript from RNases, promotes cellular uptake and endosomal escape, and contributes intrinsic adjuvant activity that favours both humoral and cell-mediated responses.^3,25,26^ The platform is also rapidly adaptable to emerging targets and is being extended to transmission-blocking veterinary vaccines.^4,30^ Whether an LNP-formulated mRNA encoding a CagW antigen can protect against *H. pylori* in vivo has, however, not been established.

Antigen choice is critical, because immunity that merely reduces colonisation is unlikely to translate into durable protection. The cag pathogenicity island (cagPAI), a ∼40-kb polymorphic locus carried by most disease-associated strains, is the dominant virulence determinant in *H. pylori* infection and differs appreciably between isolates.^15,24^ It encodes approximately 27 proteins, about 20 of which build the type IV secretion system (T4SS), the apparatus that injects the effector oncoprotein CagA into gastric epithelial cells and thereby perturbs host signalling, favours malignant transformation and raises gastric cancer risk.^24^ In *Agrobacterium tumefaciens* the homologous apparatus is the VirB/VirD4 system, assembled from eleven VirB subunits plus the coupling protein VirD4^16^, and CagA translocation by *H. pylori* depends on a set of VirB/D4-like components.^17,18^ CagW (cag9/HP0529) shares no sequence similarity with those components, yet genetic, localisation and interaction data place it as the functional counterpart of VirB6^16–18^: like VirB6 family proteins it is predicted to span the inner membrane six times, its carboxy-terminal transmembrane segment is enriched in valine, leucine and isoleucine, and it carries a tryptophan-rich motif together with a propensity for self-association.^19^ Membrane-anchored, multi-spanning and present in every cagPAI-positive strain examined, CagW is attractive as a vaccine target precisely because it is both conserved and functionally indispensable.

Antigen selection has traditionally depended on trial-and-error immunisation, but computational vaccinology now allows candidates to be triaged before any animal work begins. Immunoinformatics underpins epitope mapping, antigenicity and allergenicity screening, structural modelling and immune simulation, and reverse vaccinology exploits these tools to interrogate whole proteomes for protective candidates.^20,21^ The approach is inexpensive, scalable and safe, although its predictive value for any individual antigen is rarely checked against experimental immunogenicity. We therefore addressed two questions: whether CagW, chosen on the conservation and topology arguments set out above, behaves as a genuine protective antigen in vivo, and whether an LNP-formulated mRNA encoding it outperforms a conventional protein vaccine. CagW was first profiled computationally (antigenicity, physicochemical behaviour, secondary and tertiary structure, receptor docking and predicted B-cell epitopes); a codon-optimised CagW-Fc mRNA-LNP vaccine was then produced, characterised biophysically, and tested for immunogenicity and protective efficacy against live *H. pylori* challenge in C57BL/6 mice. To our knowledge, an mRNA-encoded CagW antigen has not previously been tested in animals.

## Methods

### Computational analyses

Sequence retrieval. The CagW amino-acid sequence of *H. pylori* strain 26695 (UniProt accession E0WID1) was downloaded from the UniProt knowledgebase (http://www.uniprot.org).

#### Antigenicity, immunogenicity, allergenicity and toxicity

The IEDB immunogenicity server (http://tools.iedb.org/immunogenicity/) was used to score the likelihood that CagW peptides would be recognised by the adaptive immune system, higher scores indicating greater predicted immunogenicity, and antigenicity was estimated with ANTIGENpro (http://tools.iedb.org/antigenpro). Sensitising potential was screened with AllerTOP v.2.0 (http://www.ddg-pharmfac.net/AllerTOP/) and AllergenFP v.1.0 (http://ddg-pharmfac.net/AllergenFP/), and toxicity with ToxinPred (http://crdd.osdd.net/raghava/toxinpred/); both AllerTOP and ToxinPred returned a negative call.

#### Physicochemical profiling and solubility

ProtParam on the ExPASy server (https://web.expasy.org/protparam/) returned the theoretical isoelectric point (pI), the estimated half-life in mammalian reticulocytes in vitro and in yeast and *E. coli* in vivo, the instability index and the aliphatic index. Solubility was predicted with Protein-Sol (https://protein-sol.manchester.ac.uk/), a sequence-based estimator that reports a score between 0 and 1, values above the 0.45 threshold indicating a readily soluble protein.

#### Secondary and three-dimensional structure

PSIPRED (http://bioinf.cs.ucl.ac.uk/psipred/) was used for secondary-structure assignment and the 3Dpro module of the Scratch Protein Predictor (uci.edu) for initial three-dimensional modelling. The crude model was refined in GalaxyRefine (https://galaxy.seoklab.org/cgi-bin/submit.cgi?type=REFINE) to improve stereochemistry and overall model quality.

#### Model validation and molecular docking

ERRAT (https://saves.mbi.ucla.edu/) scored the packing quality of the refined model, with Ramachandran statistics and a ProSA-web Z-score used as complementary checks. Coordinates for HLA-DRB1*01:01 (PDB 1AQD), TLR-2 (6NIG) and TLR-4 (4G8A) were retrieved from the Molecular Modeling Data Bank (MMDB, https://www.ncbi.nlm.nih.gov/structure/). Rigid-body docking of CagW against each receptor was performed in ZDOCK 3.0.2 and the resulting complexes rendered in PyMOL 2.5.3.

#### Conformational B-cell epitopes

Because more than 90% of B-cell epitopes are discontinuous, prediction was carried out on the folded model rather than on linear sequence. ElliPro (http://tools.iedb.org/ellipro/) was run at a threshold of 0.5 and the predicted patches were mapped onto the structure.^20,21^

#### Immune simulation

C-ImmSim (https://150.146.2.1/C-IMMSIM/index.php) was used to model the kinetics of T and B lymphocytes and of antibody production after vaccination. The default simulation volume, a random seed and the HLA allele recommended by the server were retained; each run covered 189 time steps with three injections on days 0, 14 and 28.

### Experimental validation

**Bacterial strain, cells and culture conditions.** *H. pylori* ATCC 26695 was obtained from the Henan Provincial Engineering Research Center for Industrial Microbial Strains (BNCC). Solid medium was prepared by dissolving 5.2 g of *H. pylori* agar base in 93 mL of distilled water, autoclaving at 121 °C for 15 min and, after cooling to 50–55 °C, adding 7 mL of sterile defibrinated goat blood and 1 mL of *H. pylori* supplement; the mixture was dispensed into sterile Petri dishes. Plates were incubated at 37 °C under 5% O_2_/10% CO_2_/85% N_2_ for 72–96 h.

**Plasmid linearisation and in vitro transcription.** The pIVTRup SP-CagW-Fc-Flag construct was linearised with EcoR I, purified with a standard DNA clean-up kit and checked on a 1% agarose gel, with yield and purity read on a NanoDrop 2000 (OD260 and OD260/OD280). Transcription used the HiScribe T7 ARCA mRNA Kit (NEB, USA) with N1-methylpseudouridine (m^1^Ψ) substituting for UTP. Transcript concentration was measured on the NanoDrop 2000 and integrity confirmed by 5% urea-denaturing PAGE.

**Construct design and mRNA-LNP manufacture.** A codon-optimised CagW open reading frame, preceded by a signal-peptide sequence and followed by a humanised Fc tag, was synthesised and inserted into an mRNA expression plasmid; the linearised plasmid served as template for T7-driven in vitro transcription, and purified mRNA was stored at −80 °C. For particle assembly, the ionisable lipid lipid01, cholesterol, DSPC and DMG-PEG2k were dissolved in anhydrous ethanol at a molar ratio of 20:38.5:40:1.5 to form the organic phase, and CagW-Fc mRNA was diluted in 50 mM sodium citrate buffer, pH 4, to form the aqueous phase; the two streams were combined at an N/P ratio of 5.76^22^. Mixing was performed 3:1 in a NanoAssemblr microfluidic device at 12 mL/min, and ethanol was removed by 40-fold dilution into sterile Ca^2+^- and Mg^2+^-free D-PBS, giving a final ethanol content below 1%. The formulation was concentrated to 2 mg/mL on a pre-sterilised Amicon Ultra-15 centrifugal filter (100 kDa cutoff) by centrifugation at 4,000 × g and 4 °C for 15–30 min and stored at 4 °C. Particle size and homogeneity were measured by dynamic light scattering and surface charge with a Malvern Zetasizer. Encapsulation efficiency was obtained by comparing total mRNA, released with 1% Triton X-100, with free mRNA in intact particles using the Quant-iT RiboGreen RNA Assay Kit (Thermo Fisher Scientific, USA), and morphology was examined by transmission electron microscopy.^23^

**Transfection and Western blotting.** HEK293T cells (3 × 10^6^ per dish) were plated in 10-cm dishes and maintained at 37 °C under 5% CO_2_. CagW-Fc mRNA-LNPs (20 µg) were added dropwise, mixed gently and left in contact for 24 h; untreated cells served as the negative control. Supernatants and cell lysates were harvested separately and CagW-Fc was enriched from both with a FLAG (DYKDDDDK) fusion purification kit. Aliquots containing 20 µg of protein were boiled for 10 min at 95 °C in 5× loading buffer, resolved on 10% SDS-PAGE gels and transferred onto 0.45-µm PVDF membranes by wet transfer. Membranes were blocked for 1 h in 5% skimmed milk, washed three times in PBST (0.1% Tween-20), probed with DyLight 488-conjugated goat anti-mouse IgG H&L (1:4000) for 1 h at room temperature in the dark, washed again three times in PBST and imaged.

Indirect immunofluorescence. Twenty-four hours after transfection, HEK293T cells were fixed in 4% paraformaldehyde and stained with a mouse anti-CagW polyclonal antibody (1:2000) followed by FITC-conjugated goat anti-mouse IgG (1:200). After PBS washes, nuclei were counterstained with Hoechst 33342 (1 µg/mL; Sigma) for 5 min, and green and blue signals were captured on an inverted fluorescence microscope.

**Animals and immunisation.** Specific-pathogen-free female C57BL/6 mice aged 6–8 weeks were supplied by Spearfish (Beijing) Biotechnology Co., Ltd. and housed at 22 °C and 55% ± 5% relative humidity on a 12 h light/12 h dark cycle with food and water available ad libitum. Animals were allocated at random to four groups of 17: alum alone (0.36%), luciferase mRNA-LNPs (5 µg mRNA), CagW-Fc mRNA-LNPs (5 µg mRNA) and CagW protein adsorbed to alum (80 µg). Each mouse received three intramuscular injections 14 days apart.

Anaesthesia was induced with 4–5% isoflurane in oxygen and maintained at 1.5–2% through an inhalation system, an absent paw-withdrawal reflex being taken to confirm adequate depth.

**Serum IgG, IgG1 and IgG2a by indirect ELISA.** Blood was sampled on days 14, 28 and 42. Serum (100 µL/well) was added to antigen-coated plates and bound CagW-specific IgG, IgG1 and IgG2a were detected with an HRP-conjugated goat anti-mouse secondary antibody; absorbance was read at 450 nm after washing.

**Circulating cytokines.** IL-2, IL-4, IL-12, IFN-γ and IL-17A in sera collected on days 14, 28 and 42 were quantified with commercial indirect ELISA kits (Elabscience, China) used as directed by the supplier.

**Splenocyte proliferation.** Splenic lymphocytes isolated with the Mouse Lymphocyte Isolation Medium Kit (Solarbio) were seeded at 5 × 10^5^ cells in 100 µL per well in 96-well plates and stimulated with purified CagW-Fc or CagW protein (2 µg/mL); untreated cells in DMEM and ConA-treated normal splenocytes (2 µg/mL; MCE) served as negative and positive controls, respectively. After 72 h at 37 °C in 5% CO_2_ the medium was exchanged, 10 µL of CCK-8 reagent was added per well and, after a further 4 h, absorbance was recorded at 450 nm. Every treatment was run at least in triplicate, and IL-4, IFN-γ and IL-17A in the corresponding culture supernatants were measured by ELISA.

**Flow cytometry and intracellular cytokine staining.** Splenic CD3^+^CD4^+^ and CD3^+^CD8a^+^ frequencies were determined by surface labelling: lymphocytes were incubated for 30 min at 4 °C in the dark with FITC anti-mouse CD3 (0.5 µg/10^6^ cells), APC anti-mouse CD4 (0.25 µg/10^6^ cells) and PerCP anti-mouse CD8a (0.25 µg/10^6^ cells), washed three times in pre-cooled PBS and acquired. For intracellular staining, mice were killed 14 days after the third dose and spleens were processed immediately into single-cell suspensions, washed in PBS and fractionated on the same density medium. Cells were plated at 2 × 10^6^ per well in 6-well plates and restimulated with CagW-Fc or CagW protein (2 µg/mL) for 6 h before brefeldin A was added for a further 6 h. After washing in PBS containing 0.2% BSA, cells were resuspended in 100 µL and blocked against Fc receptors with anti-CD16/CD32 for 30 min, washed three times by centrifugation and counted. Surface labelling with the same CD3/CD4/CD8a panel was repeated for 30 min at 4 °C in the dark, cells were fixed and permeabilised with the Cytofix/Cytoperm Kit (BD Biosciences, USA), and intracellular cytokines were stained for 30 min at 4 °C in the dark with PE/Cyanine7 anti-mouse IL-4 (1.0 µg/10^6^ cells), PE/Cyanine7 anti-mouse IFN-γ (1.0 µg/10^6^ cells) or PE/Cyanine7 anti-mouse IL-17A (0.25 µg/10^6^ cells). Samples were washed in pre-cooled PBS, transferred to acquisition tubes and analysed in FlowJo v10.7.1.

**Challenge.** Mice were fasted for 12 h, given 200 µL of 0.2% NaHCO_3_ by gavage and, 5 min later, inoculated with 0.5 mL of *H. pylori* suspension containing 1 × 10^9^ CFU, viability having been confirmed by plating. Survival was recorded over the following 18 days. Bacterial burdens were quantified on day 4 after challenge: gastric tissue was collected and liver was removed aseptically and homogenised with a 3-mL syringe, both tissues were plated on agar containing 5% fetal bovine serum and incubated at 37 °C in 5% CO_2_ for 48 h, and viable colonies were counted.

**Histopathology.** Mice were killed by isoflurane inhalation, and stomach, heart, liver, spleen, lung and kidney were collected from each immunisation and control group (alum and luciferase mRNA-LNP), fixed in 4% paraformaldehyde, paraffin-embedded, sectioned and stained with haematoxylin-eosin (HE).

**Statistics.** Analyses were run in Prism 8.0 (GraphPad Software). Multi-group comparisons used one-way ANOVA with Tukey’s post hoc test and two-group comparisons an unpaired Student’s t-test. Significance was set at P < 0.05 and is denoted by one (P < 0.05), two (P < 0.01) or three (P < 0.001) asterisks; ns indicates not significant.

## Results

### Predicted antigenicity, allergenicity and toxicity of CagW

The IEDB immunogenicity tool returned a score of 0.69 for CagW and ANTIGENpro returned 1.03, both well above the values normally associated with poorly immunogenic sequences. AllerTOP v.2.0, AllergenFP v.1.0 and ToxinPred each gave a negative call. Taken together, the predictions identify CagW as a strongly antigenic and immunogenic protein with no predicted sensitising or toxic liability, and therefore as a plausible *H. pylori* vaccine antigen.

### Physicochemical parameters and predicted solubility

Because physicochemical behaviour shapes both manufacturability and immunogenicity, CagW was profiled with ProtParam (Table 1). Protein-Sol returned a solubility score of 0.594, above the 0.45 cut-off, indicating that the antigen is likely to be expressed in soluble form (Figure 1A).

**Figure 1.**
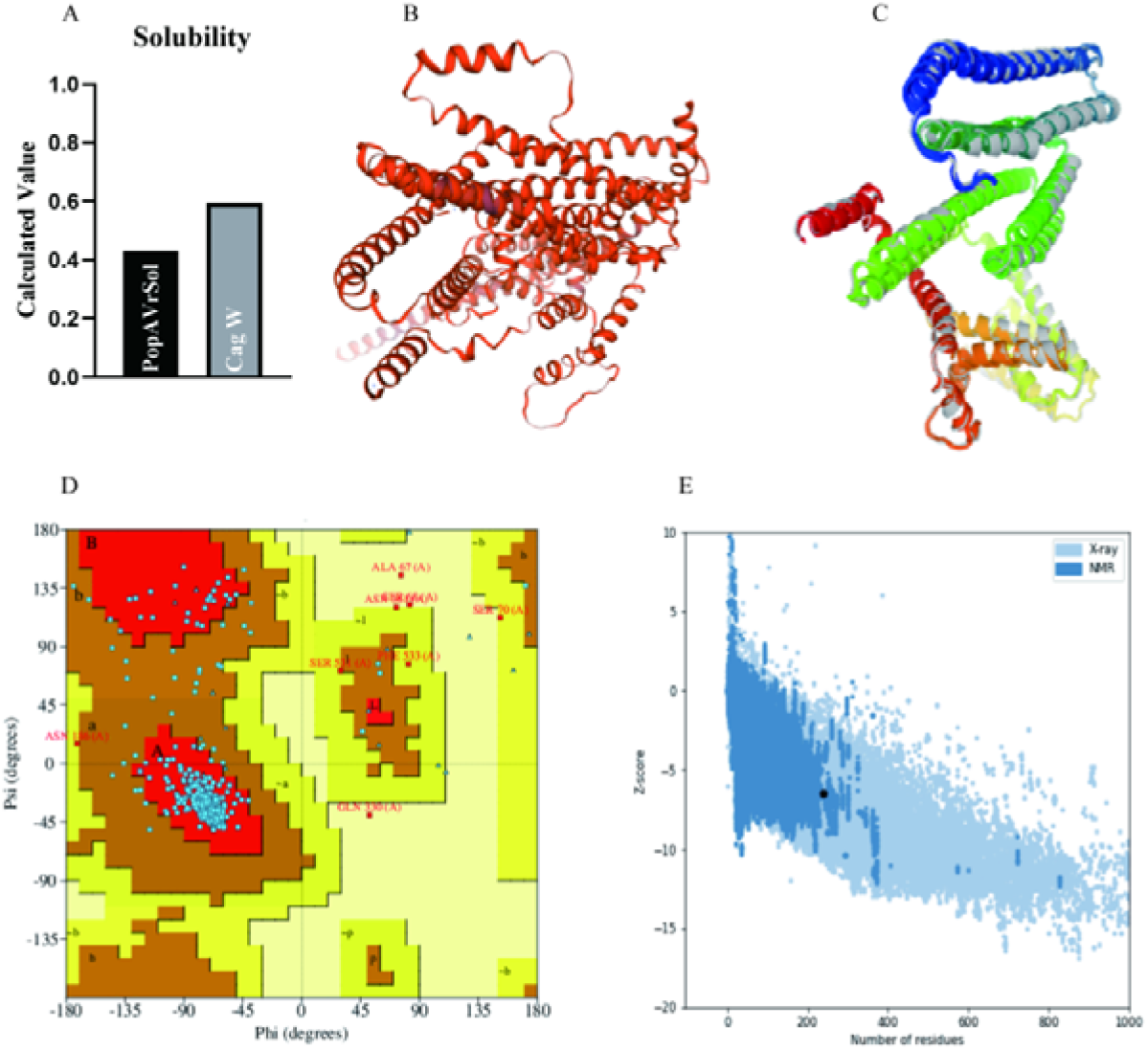
Solubility, tertiary structure and refined model of the CagW vaccine. (A) Solubility predicted by Protein-Sol. (B) Three-dimensional model built with 3Dpro before optimisation. (C) Optimised model returned by GalaxyRefine. (D) Ramachandran plot generated by PROCHECK. (E) ProSA Z-score profile.

**Table 1.** Physicochemical parameters of CagW predicted by the ExPASy ProtParam server.

| **Number of amino acids** | **Molecular weight (Da)** | **Theoretical pI** | **Estimated half-life (h)** | **Instability index** | **Aliphatic index** | **GRAVY** |
|---|---|---|---|---|---|---|
| 535 | 57,786.15 | 8.88 | 30 (mammalian reticulocytes, in vitro); >20 (yeast, in vivo); >10 (*E. coli*, in vivo) | 24.90 | 123.94 | 0.770 |

### Secondary and tertiary structure of CagW

PSIPRED assigned 58.52% of CagW residues to α-helices, 10.2% to β-strands and 31.28% to coils. An initial three-dimensional model built with 3Dpro (Figure 1B) was refined in GalaxyRefine, and the resulting models were ranked by GDT-HA, RMSD and MolProbity (Table 2). GDT-HA measures how faithfully the backbone reproduces the template geometry, RMSD quantifies deviation from the reference conformation, and MolProbity aggregates clash score, rotamer outliers and Ramachandran quality. The refined models combined high GDT-HA values with low MolProbity scores, the pattern expected of a self-consistent model (Figure 1C), and were carried forward to the analyses below.

**Table 2.** Quality metrics for the models generated by GalaxyRefine refinement.

| **Model** | **GDT-HA** | **RMSD** | **MolProbity** | **Clash score** | **Poor rotamers** | **Rama favored** |
|---|---|---|---|---|---|---|
| Initial | 1.0000 | 0.000 | 1.806 | 0.2 | 8.4 | 91.2 |
| MODEL 1 | 0.8864 | 0.573 | 1.111 | 3.2 | 0.2 | 99.1 |
| MODEL 2 | 0.8930 | 0.549 | 0.917 | 1.7 | 0.0 | 99.1 |
| MODEL 3 | 0.8818 | 0.561 | 0.986 | 2.1 | 0.0 | 99.1 |
| MODEL 4 | 0.8902 | 0.549 | 0.935 | 1.8 | 0.0 | 99.4 |
| MODEL 5 | 0.8935 | 0.539 | 1.017 | 2.4 | 0.0 | 98.9 |

### Model validation and receptor docking

ERRAT returned an overall quality factor of 90.5% for the refined model. Ramachandran analysis placed 90.5% of residues in the most favoured region, 7.8% in additionally allowed and 0.8% in generously allowed regions, so that 98.3% of residues fell within allowed conformational space (Figure 1D); ProSA-web returned a Z-score of −6.47 (Figure 1E), within the range typical of experimentally determined structures of comparable size. Docking with ZDOCK 3.0.2 produced stable complexes with HLA-DRB1*01:01 (ID 1AQD), TLR-2 (6NIG) and TLR-4 (4G8A) (Figure 2A–C), consistent with productive engagement of both antigen-presenting and innate receptors.

**Figure 2.**
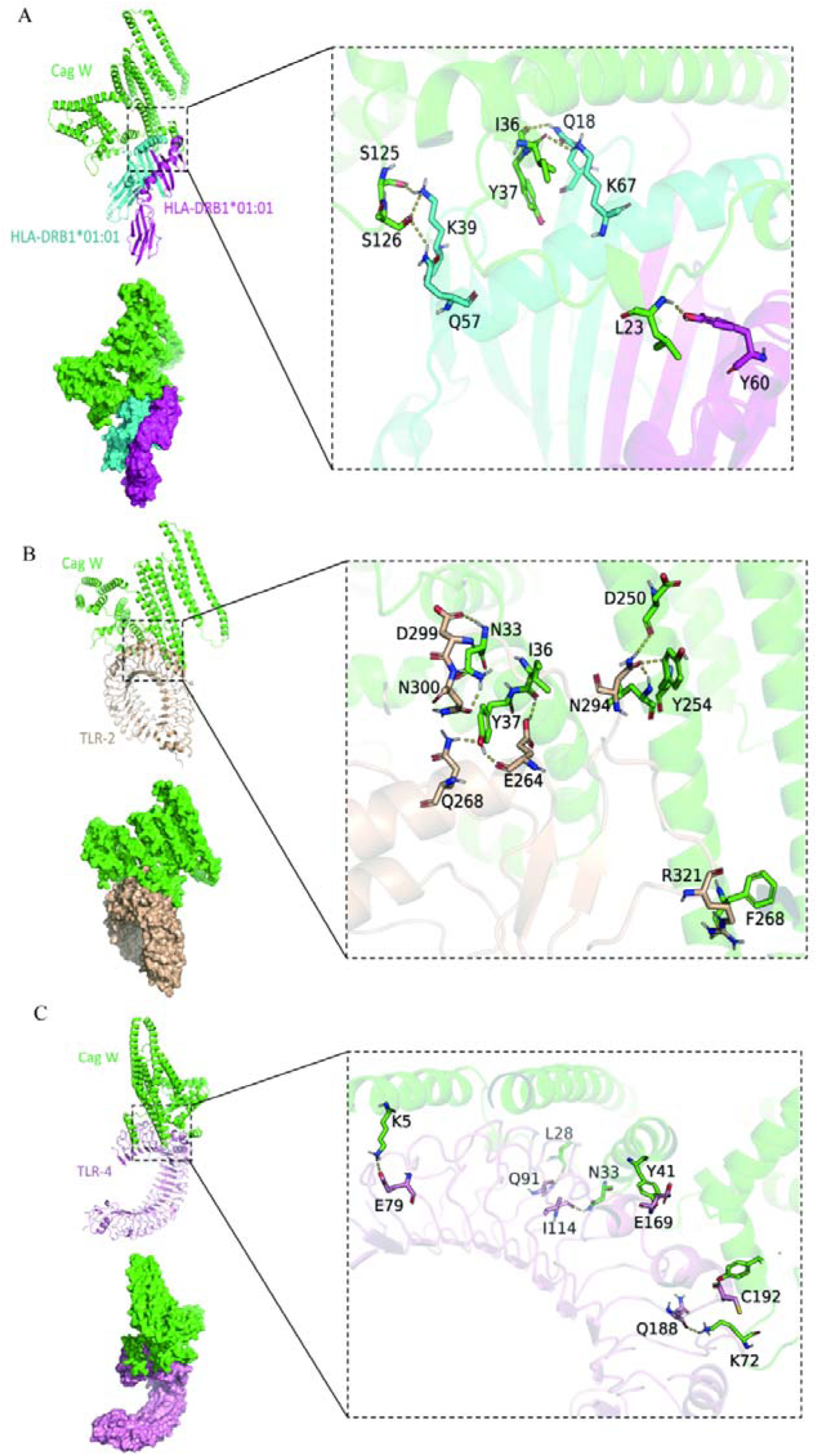
Molecular docking of CagW with HLA-DRB1*01:01, TLR-2 and TLR-4. (A) HLA-DRB1*01:01. (B) TLR-2. (C) TLR-4.

### Conformational B-cell epitopes

ElliPro identified eight conformational B-cell epitopes, with scores of 0.82 (Figure 3A), 0.742 (Figure 3B), 0.725 (Figure 3C), 0.714 (Figure 3D), 0.711 (Figure 3E), 0.646 (Figure 3F), 0.56 (Figure 3G) and 0.527 (Figure 3H). Applying a 0.69 cut-off retained six epitopes covering 227 residues (Table 3); these are distributed across both halves of the protein and are largely solvent-exposed, a topology favourable for antibody recognition.

**Figure 3.**
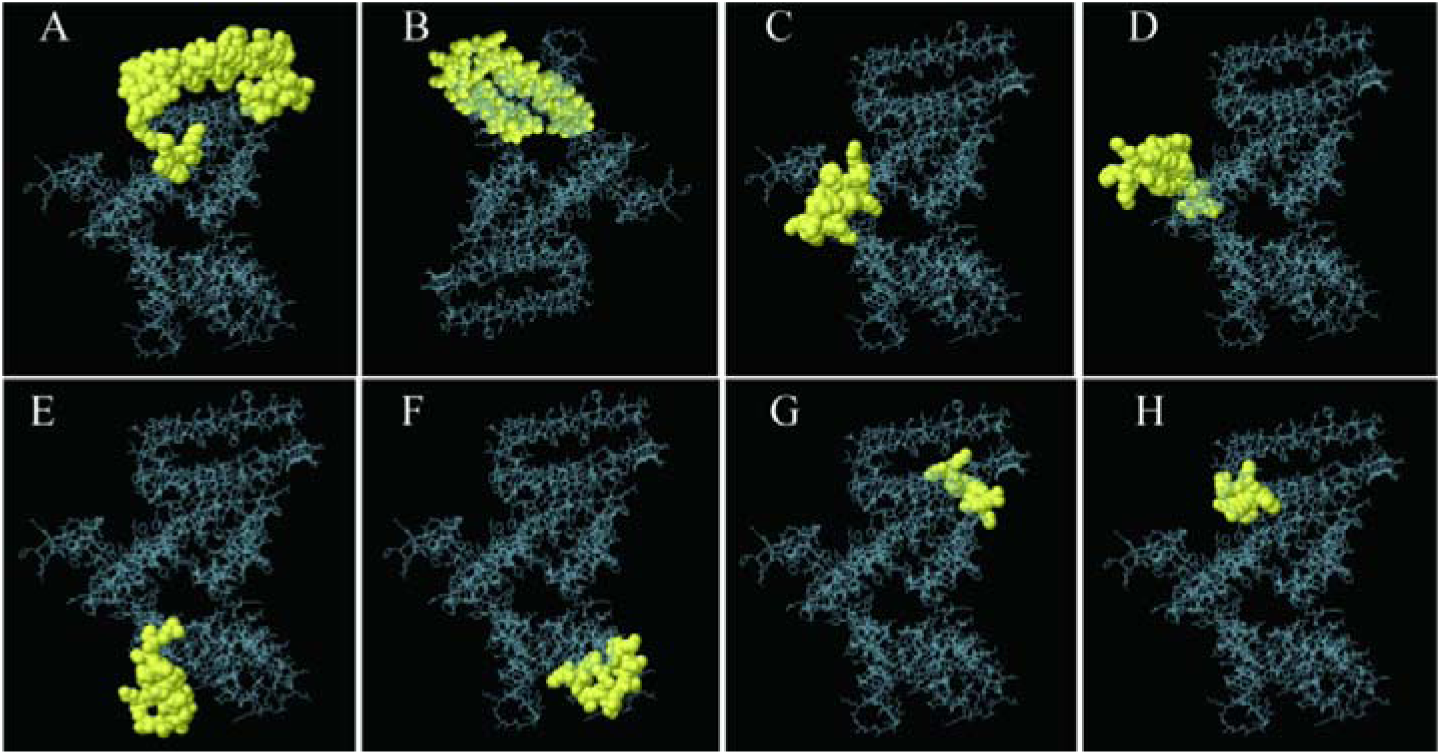
Conformational B-cell epitopes of CagW predicted by ElliPro. (A–H) Predicted epitopes are highlighted on the structure; the remaining surface is shown in grey.

**Table 3.** Conformational B-cell epitopes of CagW predicted by ElliPro.

| No. | Residues | Number of residues | Score |
|---|---|---|---|
| 1 | A:V448, A:L449, A:A450, A:L451, A:L452, A:A453, A:F454, A:N455, A:L456, A:T457, A:I458, A:T459, A:P460, A:S461, A:V462, A:A463, A:I464, A:Q465, A:N466, A:I467, A:N468, A:Q469, A:G470, A:G471, A:L472, A:G473, A:I474, A:D475, A:S476, A:T477, A:I478, A:A479, A:S480, A:L481, A:I482, A:T483, A:L484, A:F485, A:I486, A:L487, A:K488, A:G489, A:F490, A:I491, A:E492, A:T493, A:I494, A:I495, A:E496, A:S497, A:V498, A:N499, A:A500, A:I501, A:V502, A:N503, A:T504, A:I505, A:F506, A:S507, A:S508, A:V509, A:S510, A:M511, A:D512, A:G513, A:S514, A:R515, A:M516, A:D517, A:R518, A:E519, A:R520, A:D521, A:A522, A:L523, A:M524, A:I525, A:G526, A:R527, A:V528, A:G529, A:G530, A:S531, A:M532, A:F533 | 86 | 0.819 |
| 2 | A:F130, A:G131, A:W132, A:N133, A:A134, A:L135, A:I136, A:N137, A:K138, A:N139, A:I140, A:D141, A:V142, A:S143, A:M144, A:D145, A:S146, A:V147, A:L148, A:D149, A:N150, A:L151, A:S152, A:N153, A:K154, A:I155, A:Q156, A:N157, A:F158, A:T159, A:K160, A:G161, A:G162, A:V163, A:E164, A:D165, A:N166, A:V167, A:K168, A:G169, A:N170, A:I171, A:L172, A:L173, A:Q174, A:I175, A:I176, A:G177, A:S178, A:I179, A:T180, A:A181, A:Q182, A:A183, A:S184, A:T185, A:N186, A:I187, A:T188, A:E199 | 60 | 0.746 |
| 3 | A:F265, A:I266, A:S267, A:F268, A:M269, A:K270, A:L271, A:R272, A:D273, A:I274, A:S275, A:N276, A:G277, A:I278, A:Q279, A:I280, A:N281, A:I282, A:G283, A:F284, A:Y286, A:M287 | 22 | 0.738 |
| 4 | A:L50, A:T51, A:N52, A:S54, A:F55, A:A56, A:D57, A:I58, A:L59, A:T60, A:G61, A:Y62, A:T63, A:S64, A:I65, A:G66, A:A67, A:S68, A:G69, A:S70, A:G71, A:K72, A:S73, A:S74, A:G75, A:Q76, A:G77 | 27 | 0.71 |
| 5 | A:F2, A:N3, A:I4, A:K5, A:T7, A:F8, A:L9, A:V10, A:T11, A:I12, A:I13, A:S14, A:L17, A:I18, A:V19, A:P20, A:N21, A:W22, A:L23, A:K24, A:A25, A:I26, A:D27, A:L28, A:P29, A:I30, A:V31 | 27 | 0.707 |
| 6 | A:T201, A:A202, A:N203, A:K204, A:Q206 | 5 | 0.701 |

### Simulated innate and adaptive responses to CagW vaccination

C-ImmSim predicts how human immune-cell populations evolve after vaccination. Dendritic cells (DCs), the most efficient antigen-presenting cells (APCs), were markedly activated, total DC numbers stabilising near 200 cells/mm^3^ after stimulation (Figure 4A). Macrophage activation followed the same course, remaining at approximately 196 cells/mm^3^ across the three injections (Figure 4B).

**Figure 4.**
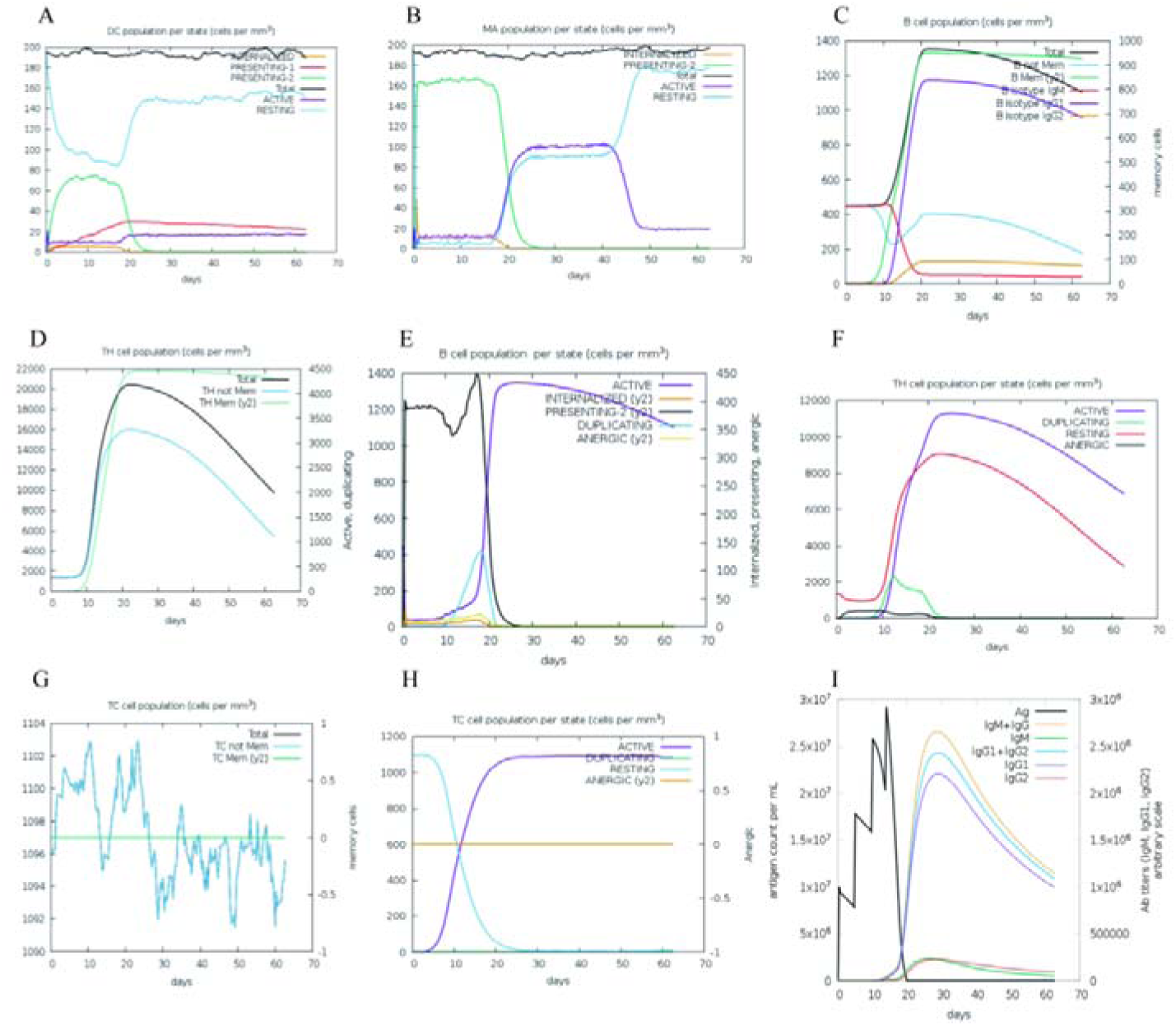
Immune simulation of innate and adaptive responses using C-ImmSim. (A) Dendritic-cell numbers and status. (B) Macrophage numbers and status. (C) B-cell populations. (D) Helper T-cell populations. (E) Helper T-cell status. (F) Cytotoxic T-cell population. (G) Cytotoxic T-cell status. (H) Predicted CagW-specific IgG, IgM, IgG1 and IgG2 levels.

B cells, CD4^+^ T cells and CD8^+^ T cells form the adaptive arm relevant to *H. pylori* clearance, and antibody-mediated mechanisms are now recognised as more than ancillary. Simulated B-cell numbers in every state rose to a maximum of about 1,400 cells/mm^3^ after the third injection (Figure 4C). Three successive peaks were evident for helper T cells, with non-memory, memory and total TH populations reaching 16,000, 22,000 and 21,000 cells/mm^3^ after the third dose (Figure 4D), while activated and resting CD4^+^ T cells peaked at approximately 12,000 and 9,000 cells/mm^3^ (Figure 4E).

CD8^+^ T cells contribute to bacterial clearance through perforin and granzyme B. Non-memory cytotoxic T lymphocytes (CTLs) reached 1,104 cells/mm^3^ after the third stimulation (Figure 4F), and activated TC cells peaked at 1,100 cells/mm^3^ over the same interval (Figure 4G).

### Simulated antibody response to CagW vaccination

Antibody titres rose steeply in the simulated CagW group. IgM peaked at 2.5 × 10^6^/mL after the third injection and then declined gradually, whereas combined IgG1 plus IgG2 peaked at 2.5 × 10^7^/mL and decayed similarly; individual maxima were 2.3 × 10^7^/mL for IgG1 and 2.7 × 10^7^/mL for IgG2 (Figure 4H). **Construction and biophysical characterisation of CagW-Fc mRNA-LNPs**

Double digestion of the CagW recombinant plasmid with EcoR I and Bgl II released the expected ∼2,454-bp SP-CagW-Fc-Flag fragment (Figure 5A), confirming correct assembly of the construct.

**Figure 5.**
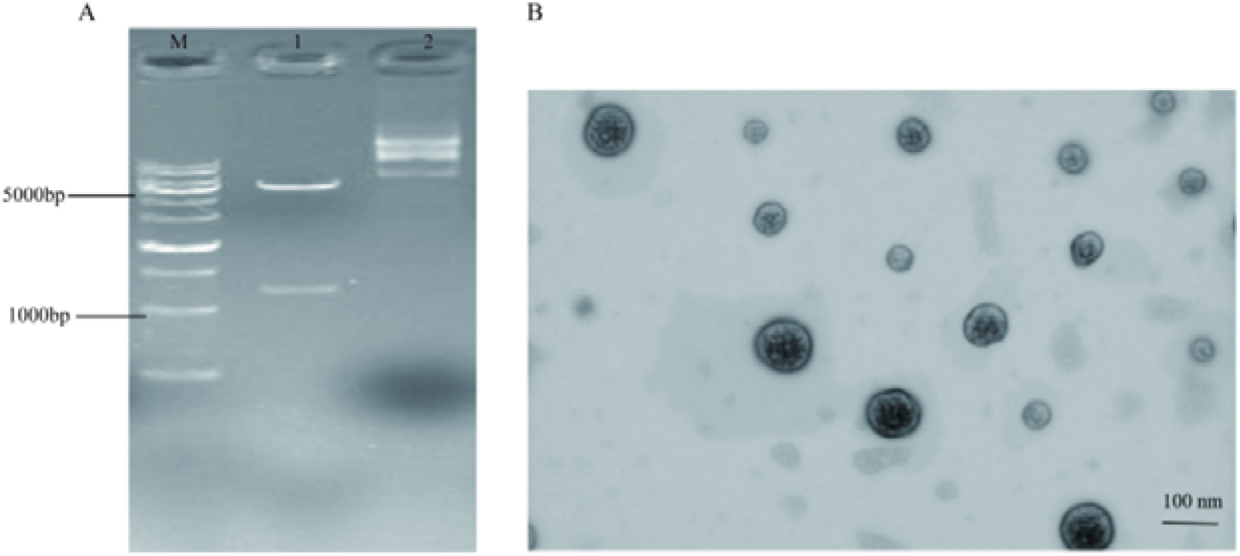
Recombinant plasmid digestion and particle morphology. (A) Double digestion of the SP-CagW-Fc-Flag plasmid on a 1% agarose gel. (B) Transmission electron micrograph of mRNA-LNPs; scale bar, 100 nm.

Because CagW is an inner-membrane protein whose folding in bacteria depends on accessory cag gene products, expression in a mammalian host was expected to be difficult, and the mRNA format was chosen partly to bypass this constraint. The construct carries a Cap0 structure, a human signal peptide that routes the antigen into the secretory pathway and a human Fc tag for detection and multimerisation (Figure 5). Transcripts were generated by T7-mediated in vitro transcription with complete replacement of uridine by m^1^Ψ and assembled by microfluidics into LNPs composed of lipid01, DSPC, cholesterol and DMG-PEG2k at a molar ratio of 20:40:38.5:1.5. Dynamic light scattering gave a mean diameter of 93.2 nm and a polydispersity index of approximately 0.13, encapsulation efficiency was about 90% by the RiboGreen assay, and the mean surface charge at physiological pH was −4.815 mV. Negatively stained particles appeared as uniform solid spheres (Figure 5B).

### Antigen expression in HEK293T cells

IFA confirmed translation of the mRNA: HEK293T cells exposed to CagW-Fc mRNA-LNPs emitted green fluorescence, whereas untreated cells remained negative, in line with earlier work in which a mouse polyclonal anti-CagW antibody detected antigen in transfected cells (Figure 6A). Western blotting with an anti-FLAG antibody revealed stronger CagW-Fc bands in transfected than in control cultures (Figure 6B and C).

**Figure 6.**
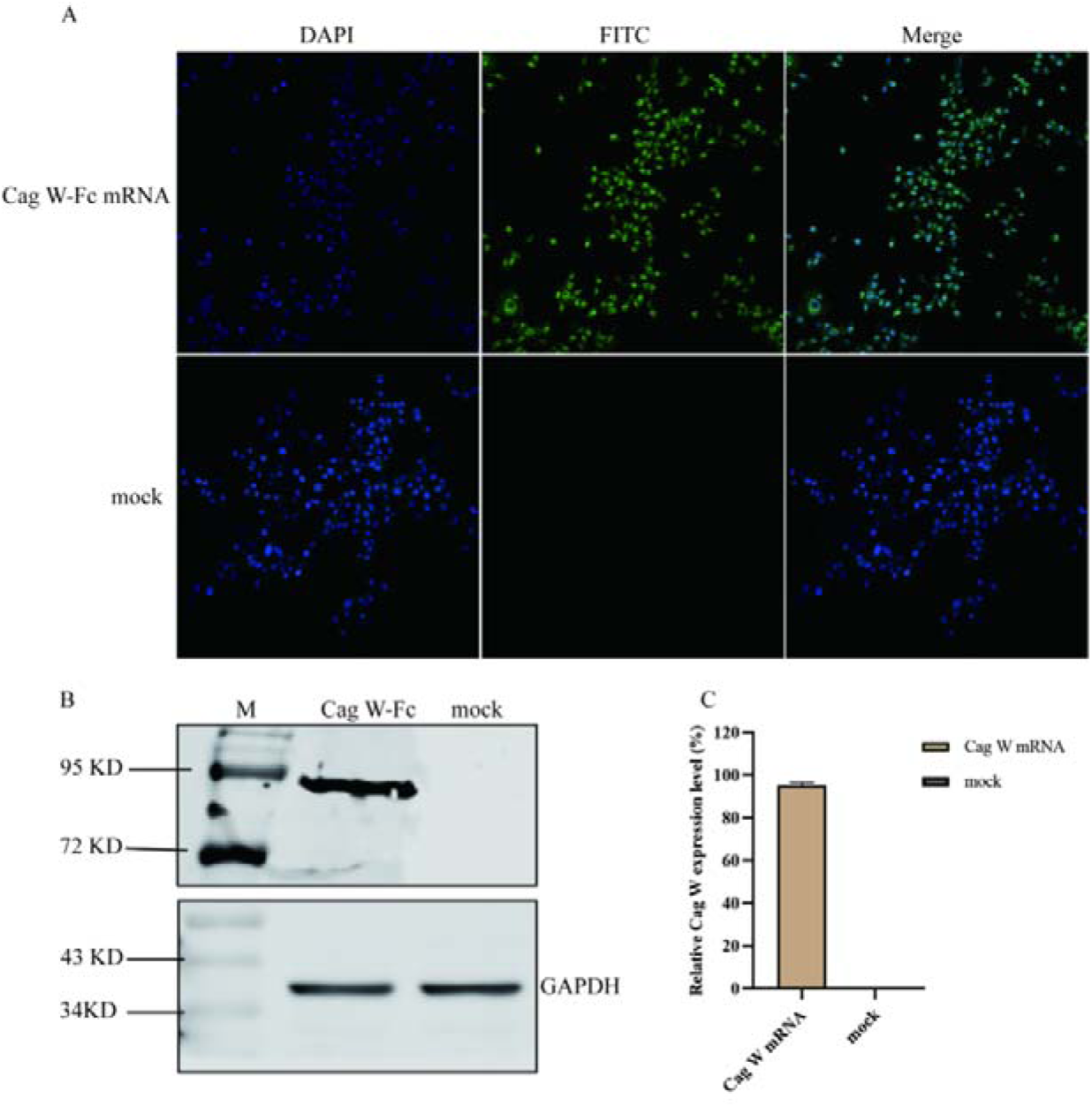
Antigen expression in vitro. (A) HEK293T cells treated for 24 h with mRNA-LNPs and examined by IFA; the secondary antibody was labelled with green fluorescence and nuclei were counterstained with DAPI (blue). (B) Western blot with an anti-FLAG primary antibody. (C) Quantification of CagW-Fc expression.

### Immunogenicity and humoral responses to CagW-Fc mRNA-LNPs

Both vaccine formats were then compared in mice. No overt change in appearance or behaviour followed any injection. Serum was screened by ELISA: CagW-specific IgG was already detectable on day 14 in the CagW-Fc mRNA-LNP and CagW protein groups, rose steadily thereafter and peaked on day 42, whereas the alum and luciferase mRNA-LNP groups remained at baseline (Figure 7A–C). Titres in both vaccine groups were significantly higher than in either control (P < 0.001), confirming that the mRNA-LNP formulation is immunogenic and drives antigen-specific IgG production.

**Figure 7.**
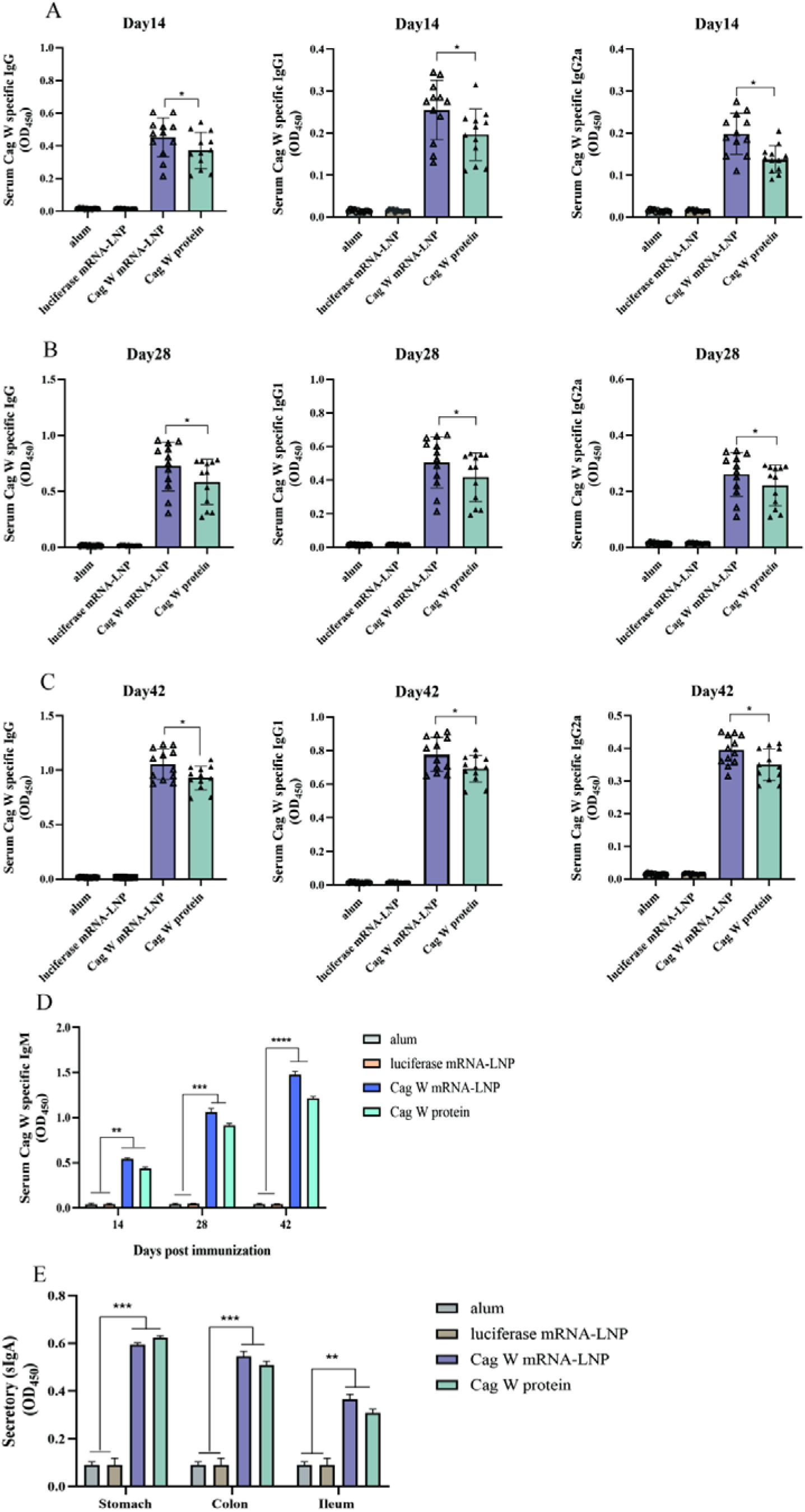
Serum CagW-specific antibody levels measured by ELISA. (A–C) IgG, IgG1 and IgG2a. (D) IgM. (E) sIgA in stomach, colon and ileum. (F) IgA. (G) Gastric mucosal sIgA.

IgM, a potent agglutinating and bactericidal immunoglobulin that dominates the earliest phase of a humoral response, was significantly higher in the CagW mRNA and CagW protein groups than in the alum and luciferase mRNA-LNP groups by day 28 and peaked on day 42 (Figure 7D), an early line of defence likely to matter in the first days after exposure.

Secretory IgA and IgA followed the same course: both were significantly elevated in the two vaccine groups by day 28 and maximal on day 42 relative to the controls (Figure 7E and F), and gastric mucosal sIgA titres were likewise significantly higher in vaccinated animals (Figure 7G).

Circulating IL-2, IL-4, IL-12, IFN-γ and IL-17A were measured by ELISA. All five were significantly higher in the CagW mRNA and CagW protein groups than in the alum and luciferase mRNA-LNP groups (Figure 8A–E), the difference in IFN-γ, IL-17A, IL-4 and IL-12 being evident from day 14 through day 42, whereas neither control group showed any shift in these analytes. A response spanning Th1 (IFN-γ, IL-12), Th2 (IL-4) and Th17 (IL-17A) axes implies that LNPs contribute adjuvant activity beyond simple delivery. Antigen-specific recall was tested by restimulating splenocytes with recombinant eukaryotic CagW-Fc or CagW protein (2 µg/mL). The stimulation index was significantly higher in CagW mRNA-immunised mice than in any control group (Figure 9A). IFN-γ, IL-4 and IL-17A released into the culture supernatant after restimulation were likewise significantly higher in this group (Figure 9B–D), with IFN-γ and IL-4 dominating and IL-17A comparatively modest, the profile expected of a CagW-specific memory T-cell response.

**Figure 8.**
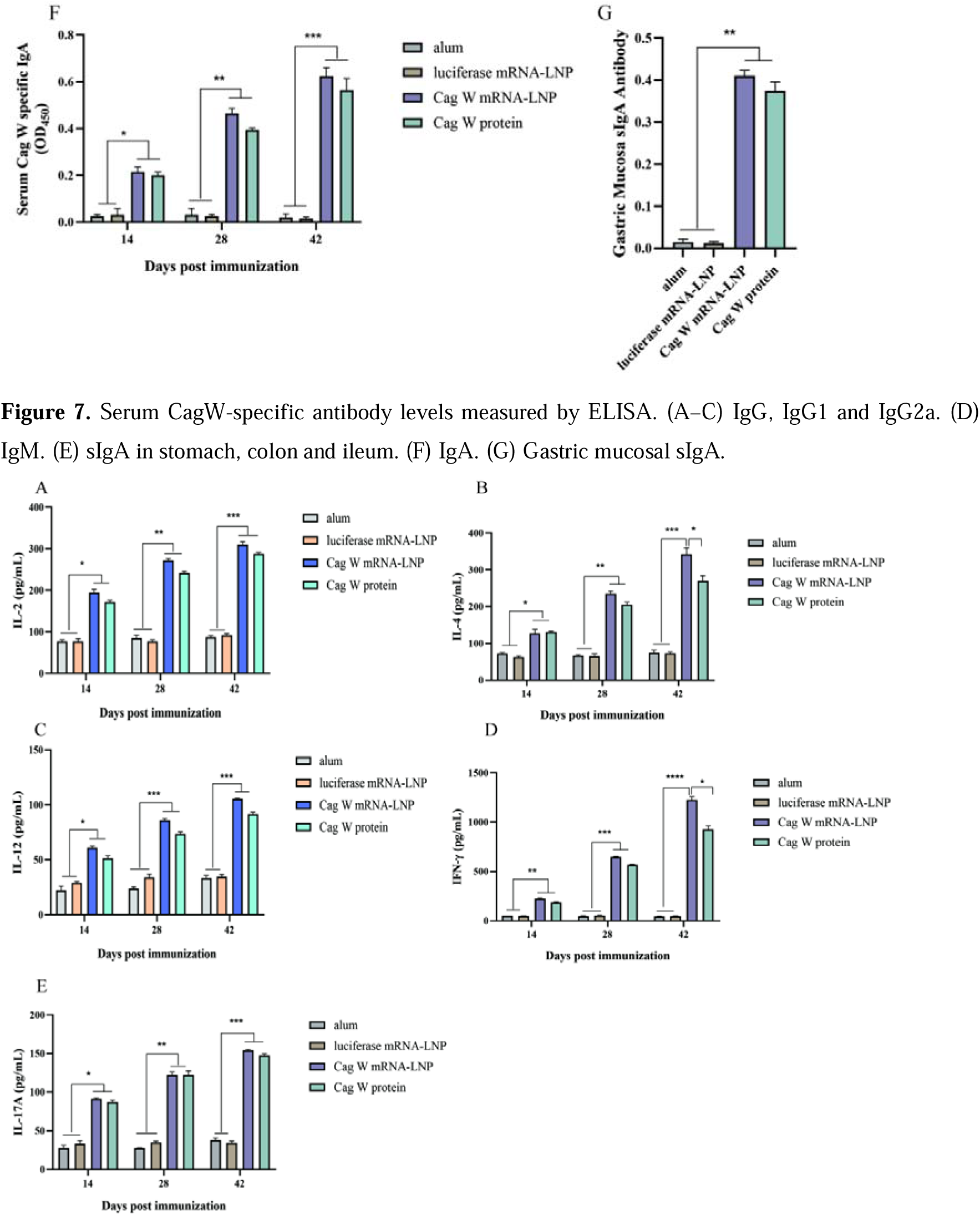
Serum cytokine levels measured by ELISA. (A) IL-2. (B) IL-4. (C) IL-12. (D) IFN-_γ_. (E) IL-17A.

**Figure 9.**
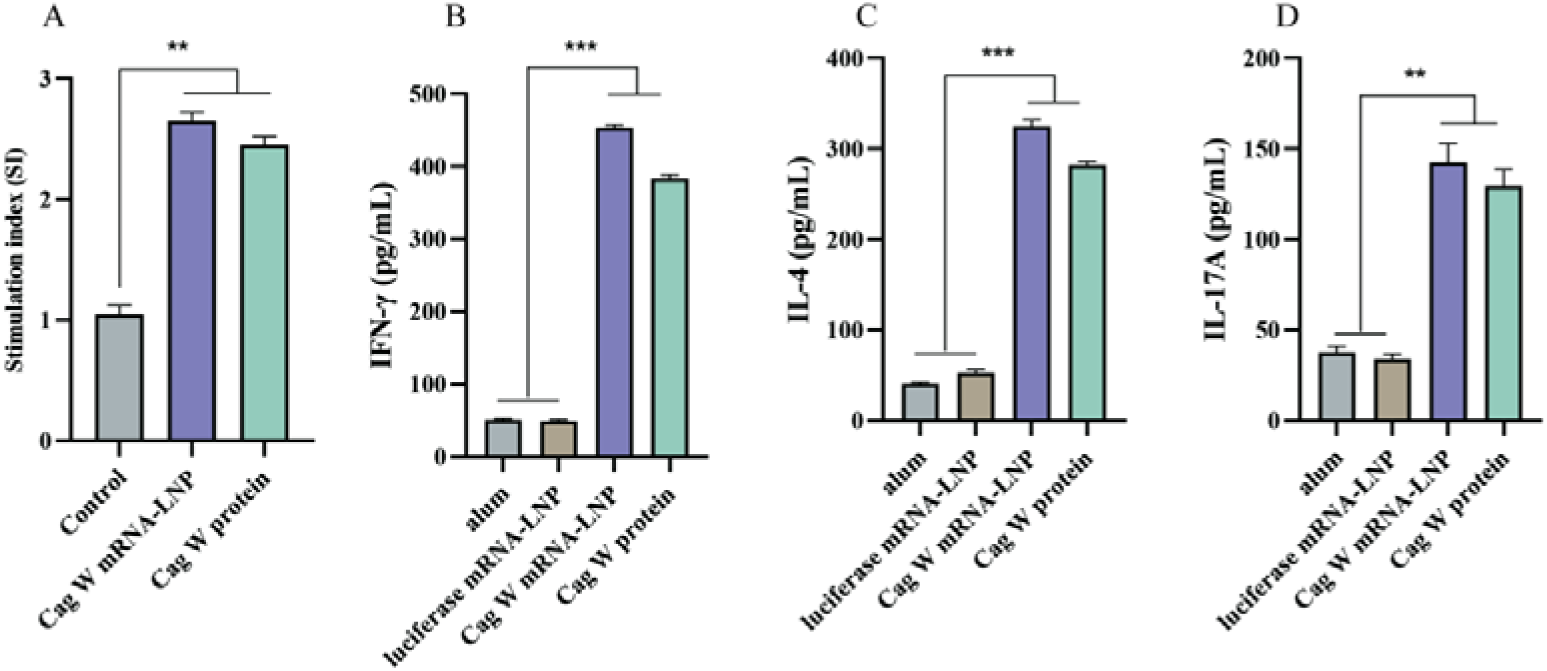
CagW-specific T-cell responses. (A) Lymphocyte proliferation after 72 h of stimulation with CagW protein, measured by CCK-8; the stimulation index is the ratio of the mean OD of stimulated to unstimulated wells. (B–D) IFN-_γ_, IL-4 and IL-17A released by restimulated lymphocytes, measured by ELISA.

### T-cell subsets and intracellular cytokines after mRNA-LNP vaccination

To resolve the cellular response further, splenic CD4^+^ and CD8^+^ frequencies were enumerated by flow cytometry 42 days after the first dose. Both subsets were significantly expanded in the mRNA-LNP group relative to the controls (Figure 10A and B), and T-cell proliferation was significantly greater than in the protein vaccine group.

**Figure 10.**
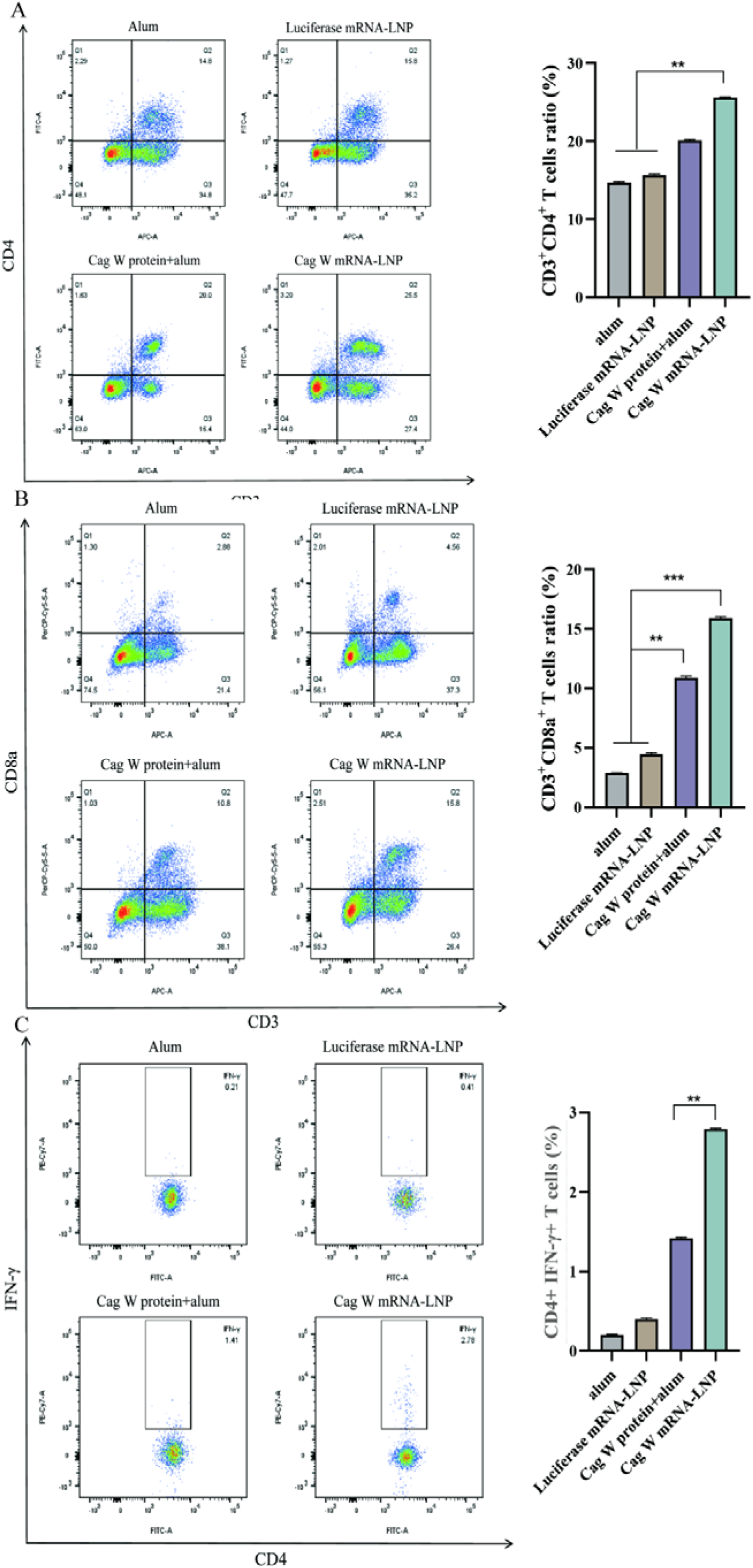
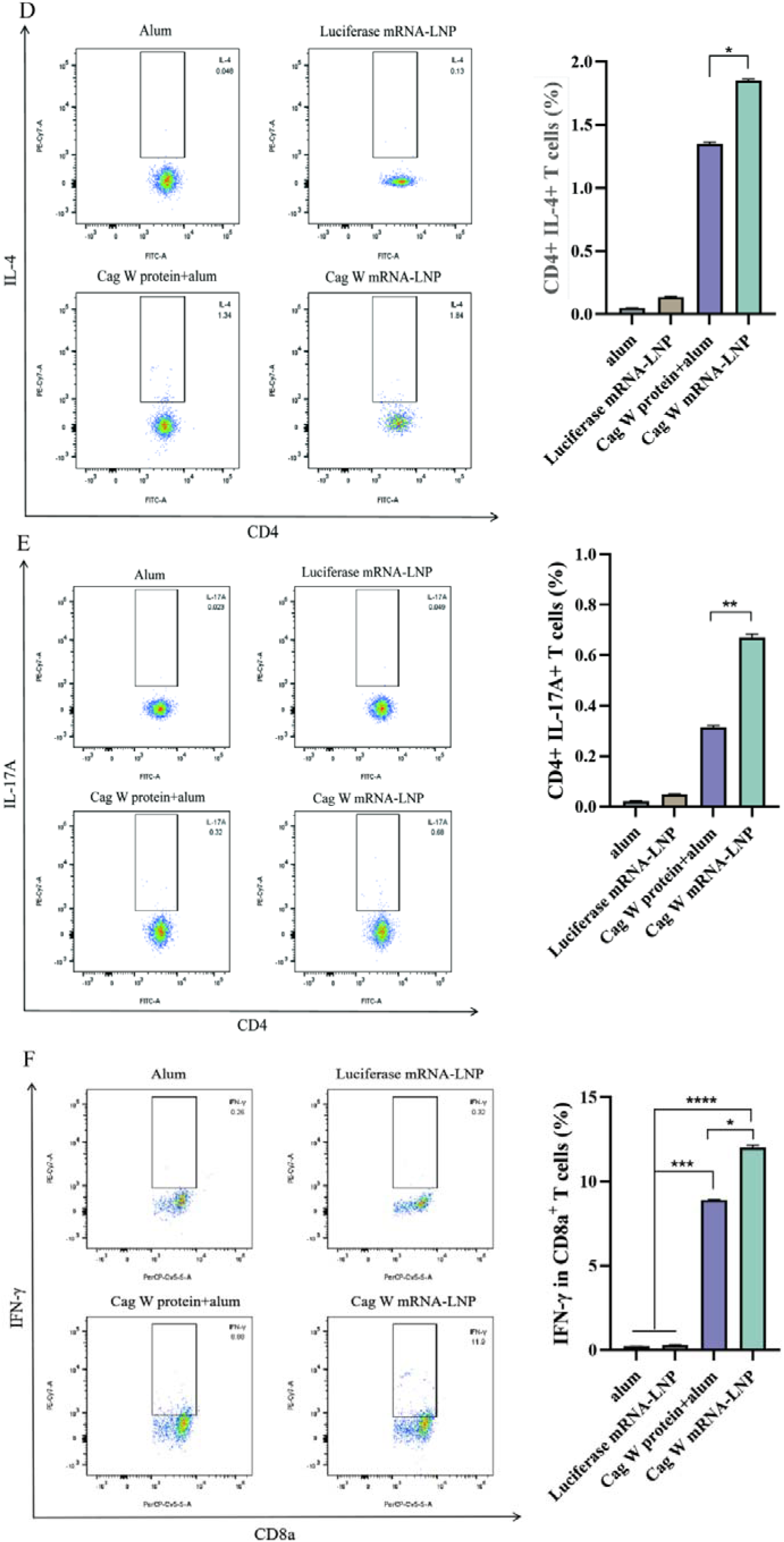
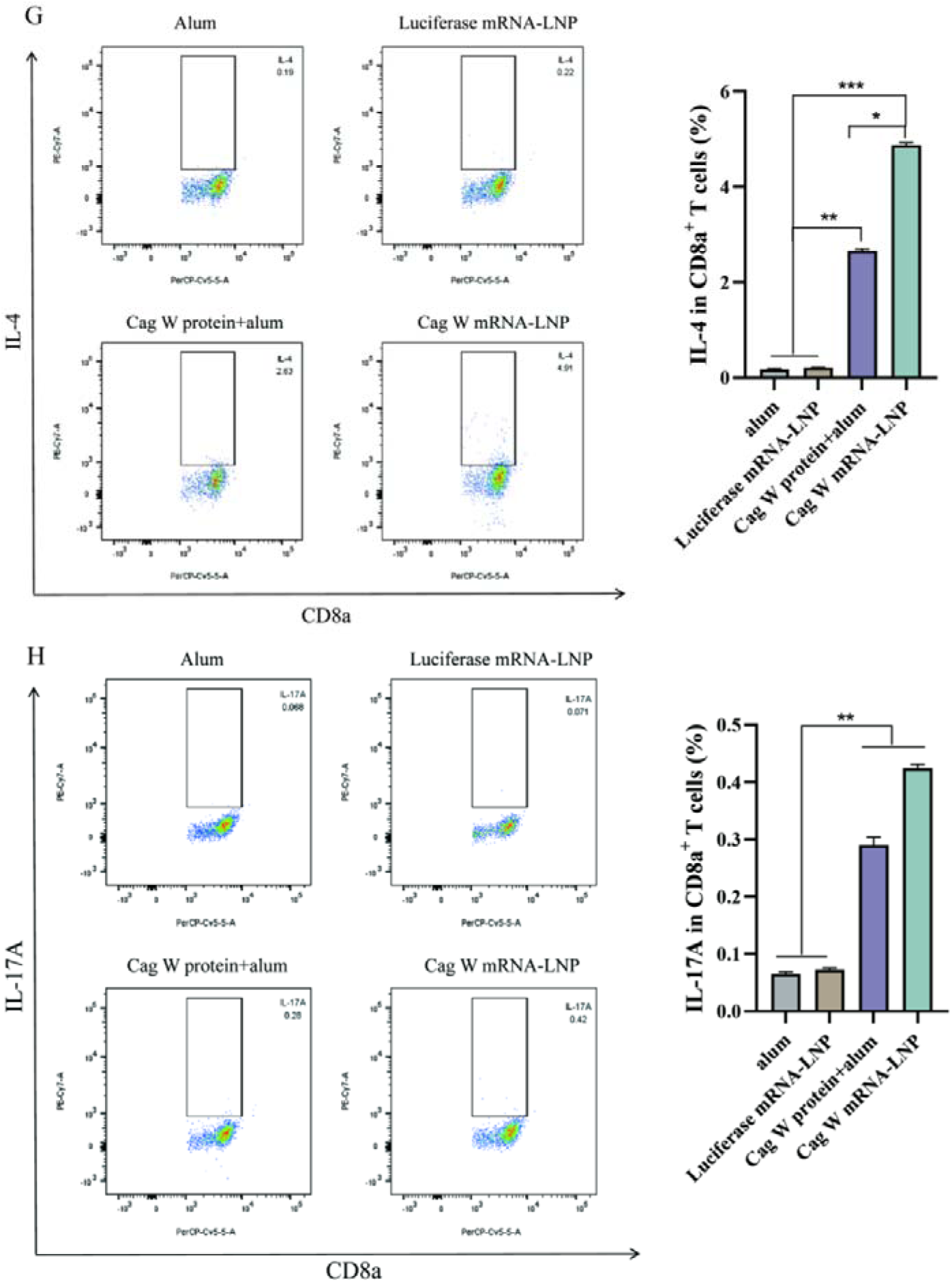
T-cell responses and cytokine expression induced by the mRNA-LNPs. (A) CD3^+^CD4^+^ T cells. (B) CD3^+^CD8a^+^ T cells. (C) IFN-_γ_^+^CD4^+^ cells. (D) IL-4^+^CD4^+^ cells. (E) IL-17A^+^CD4^+^ cells. (F) IFN-_γ_^+^ CD8a^+^ cells. (G) IL-4^+^ CD8a^+^ cells. (H) IL-17A^+^ CD8a^+^ cells.

Intracellular staining showed significantly more IFN-γ^+^ and IL-4^+^ T cells in the mRNA-LNP group at the same time point (Figure 10C–H), whereas IL-17A^+^ T-cell frequencies were unchanged.

### CagW-Fc mRNA-LNPs protect against live H. pylori challenge

Protective efficacy was assessed by gavage challenge with 1 × 10^9^ CFU of *H. pylori* ATCC 26695, the alum and luciferase mRNA-LNP groups serving as negative controls and the CagW protein group as a positive control. Bacterial recovery from liver and stomach was lower in both vaccine groups than in the controls (Figure 11A–C), and urease activity in gastric tissue was significantly reduced (Figure 11D). The CagW-Fc mRNA-LNP vaccine therefore conferred substantial protection against infection.

**Figure 11.**
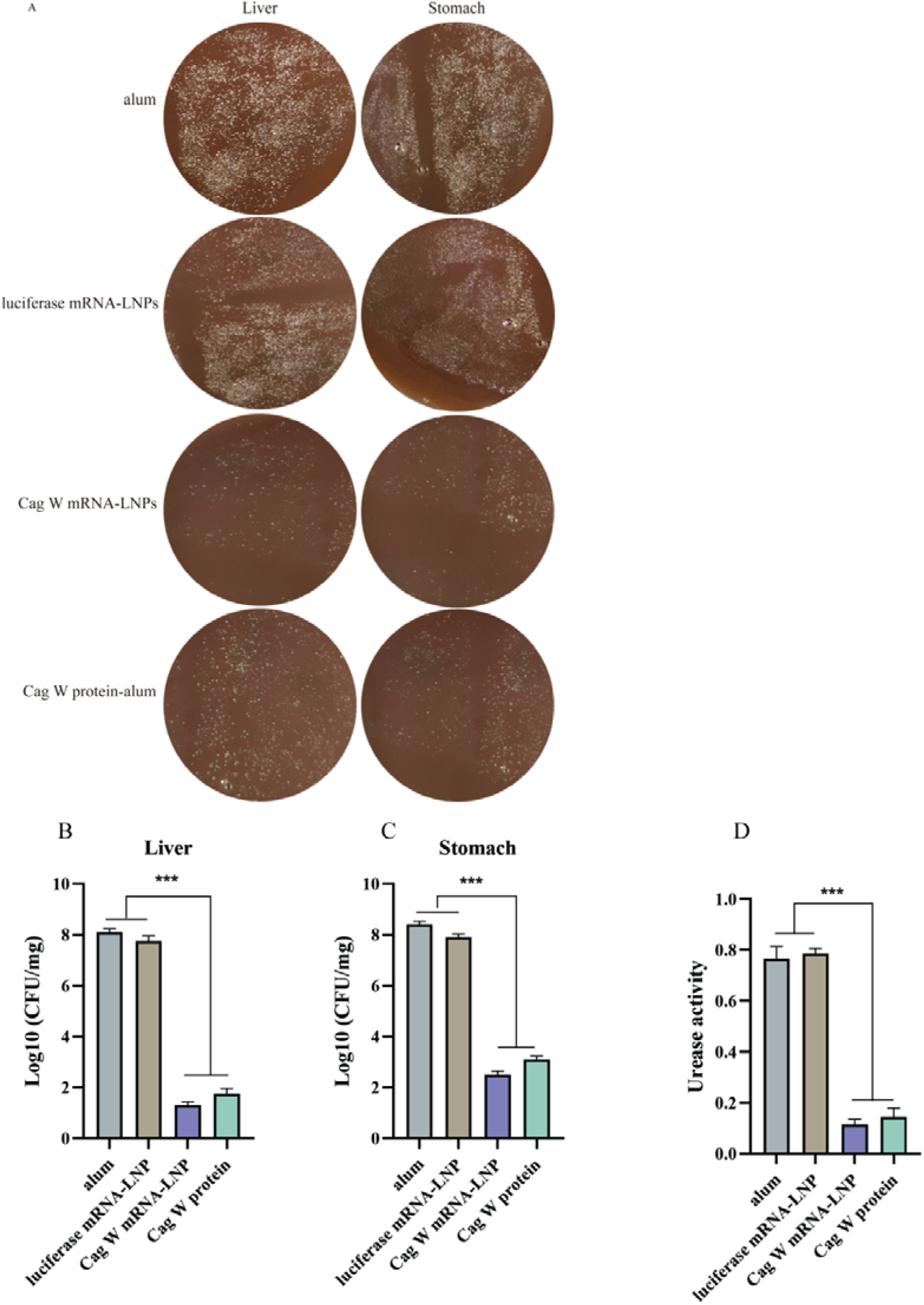
Residual bacterial counts and urease activity after *H. pylori* infection. (A) Cultures from liver and stomach suspensions. (B) Residual CFU in liver. (C) Residual CFU in stomach. (D) Urease activity in gastric tissue.

### Survival and body-weight changes after challenge

Survival was monitored for 18 days after challenge (design summarised in Figure 12A). No mouse in the alum or luciferase mRNA-LNP groups survived beyond day 4, whereas cumulative survival reached 77.8% in the CagW protein group and 100% in the CagW-Fc mRNA-LNP group (Figure 12B). Body weight fell sharply in both control groups after challenge but was essentially unchanged in the two vaccine groups (Figure 12C).

**Figure 12.**
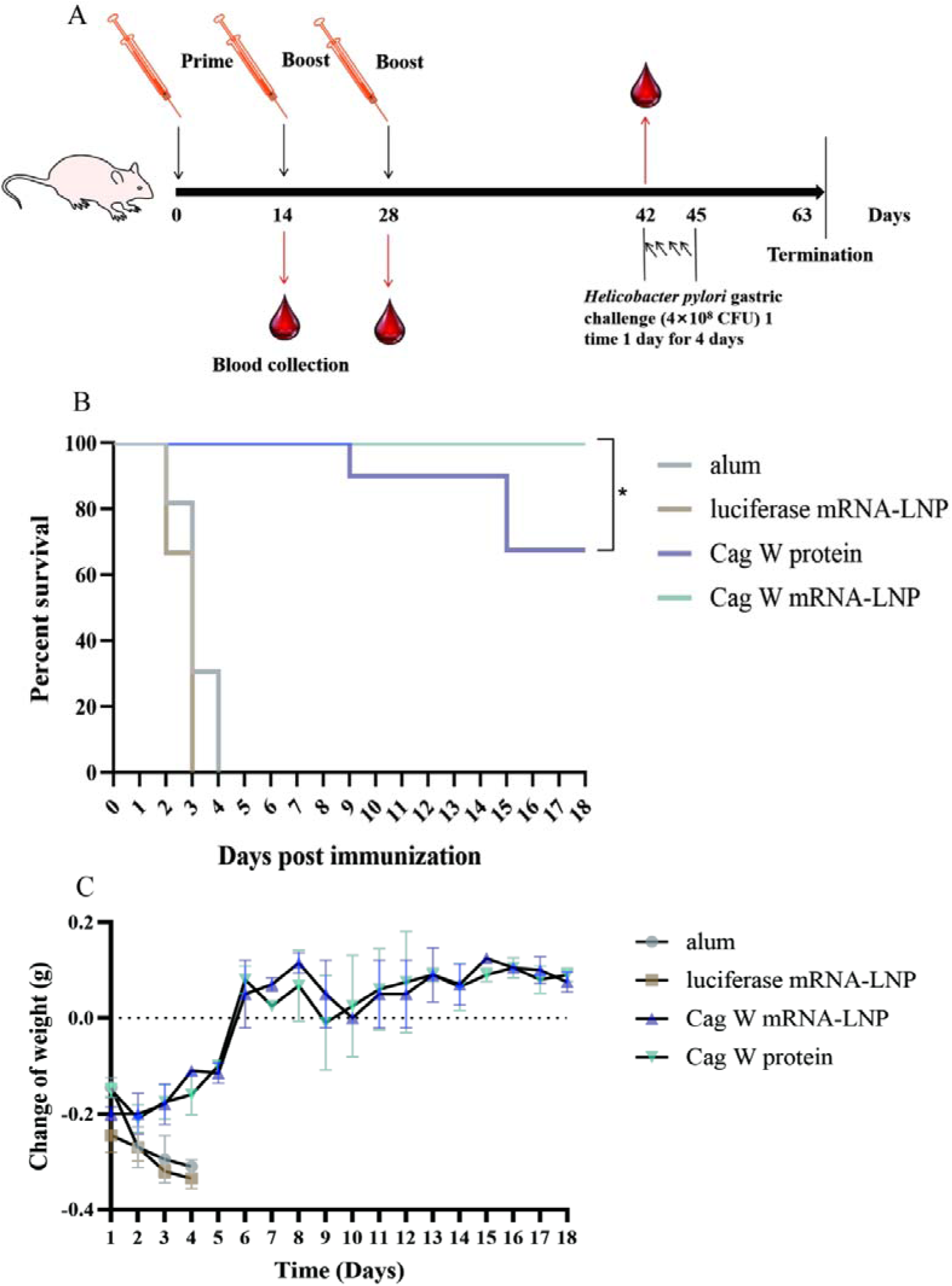
Survival and body-weight change after challenge. (A) Schematic of the immunisation and challenge schedule. (B) Survival curves. (C) Body-weight curves over the 18 days after challenge.

### Histopathology after challenge

HE-stained sections from the alum and luciferase mRNA-LNP groups showed necrotic lesions in stomach, heart, lung, liver, spleen and kidney. Animals vaccinated with CagW-Fc mRNA-LNPs or CagW protein, by contrast, showed far less infection-related damage, with efficient clearance of invading bacteria and preserved tissue architecture in the stomach and other organs (Figure 13).

**Figure 13.**
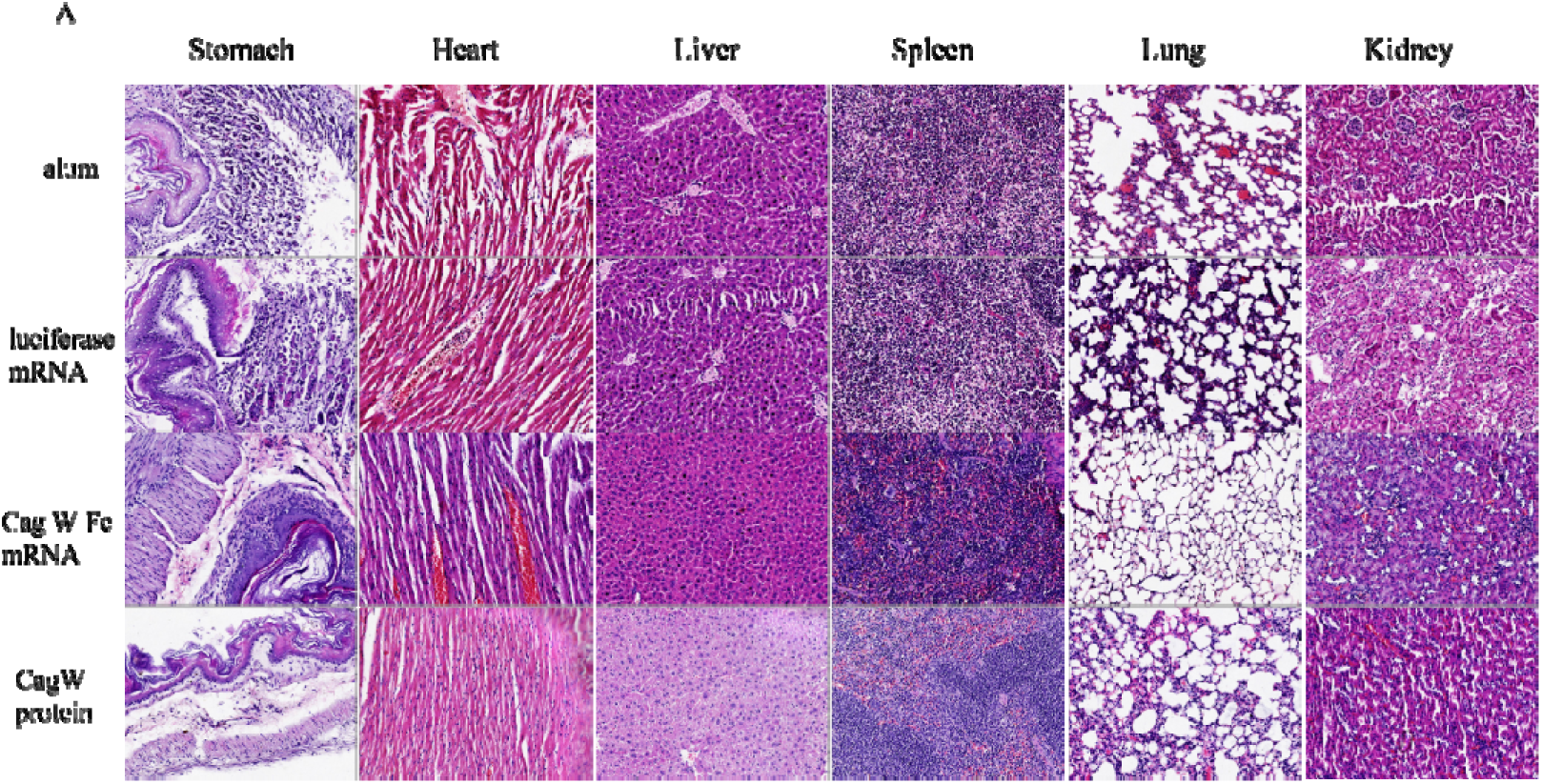
Histological safety assessment of vaccinated mice. HE staining; scale bar = 100 µm.

## Discussion

Eradication therapy for *H. pylori* is becoming less reliable as multidrug resistance spreads, and the pipeline of new antimicrobials is thin; mobilising protective immunity is therefore an increasingly attractive alternative.^25^ Progress has been slowed by the sheer complexity of *H. pylori* pathogenesis, in which several virulence factors act in concert, and by the absence of a formulation that achieves reproducible sterile immunity. Here we show that a CagW-encoding mRNA packaged into LNPs by microfluidic mixing elicits both arms of adaptive immunity and protects mice against live challenge. Two features of the platform deserve emphasis. First, LNPs are non-viral vectors with intrinsic adjuvant activity that have already been used to deliver nucleic acids and, when coupled to monoclonal antibodies, chemotherapeutics.^26^ Second, LNP-based delivery has repeatedly been shown to broaden and balance Th1/Th2 responses.^25,26^ Particle size is one determinant of that effect: small LNPs are taken up by alveolar macrophages, which release IL-12, TNF-α and IL-18 and thereby drive IFN-γ production.^26–28^ In the present study the CagW-Fc mRNA-LNPs outperformed both control formulations in lymphocyte proliferation and cytokine output, consistent with a size- and composition-dependent adjuvant effect.^29,30^ We therefore regard the LNP as an active component of this vaccine rather than an inert carrier, a view supported by the protection achieved with mRNA-LNPs against other bacterial and viral pathogens.^23,31^ CagW was selected on structural and functional grounds and then validated experimentally. The predicted secondary-structure composition (58.5% α-helix, 10.2% β-strand, 31.3% coil) leaves much of the surface solvent-exposed, and ElliPro mapped several accessible patches that could serve as antibody footprints; the antigenicity score of 1.03 is correspondingly high. Immunisation with either the mRNA-LNP or the protein vaccine raised CagW-specific IgG, IgG1, IgG2a and IgM substantially, and antibody-mediated mechanisms are known to contribute to clearance of this organism.

Encapsulation served three purposes here: it shielded the transcript, it prolonged antigen exposure, and it preserved biological activity through manufacture. The particles were smooth, evenly dispersed, non-aggregated and spherical, and transfection of HEK293T cells confirmed that the encapsulated mRNA was translated into the intended antigen. Comparable slow-release behaviour has been described for other LNP-formulated mRNA vaccines, and our physical and functional data indicate that neither the manufacturing process nor particle formation compromises potency.

Protection against *H. pylori* is usually attributed to Th1-polarised responses, although antibody and Th17 contributions are increasingly recognised. In our comparison, CagW mRNA-LNPs produced stronger splenocyte proliferation than either control and significantly increased IL-2, IL-4, IL-12, IFN-γ and IL-17A relative to all other groups, accompanied by expansion of CD4^+^ and CD8^+^ subsets and of IFN-γ^+^ and IL-4^+^ cells. This combination of humoral and cell-mediated effector mechanisms is a plausible explanation for the reduction in gastric bacterial load, the fall in urease activity and the survival benefit observed after challenge. The finding that LNPs promote dendritic-cell maturation through interferon signalling offers a mechanistic bridge between the formulation and the cellular responses measured here.

Several caveats should be borne in mind. A single mouse strain and a single challenge dose were used, protection was not benchmarked against adjuvanted Th1-driving formulations, and the correlates of protection were inferred rather than established formally by passive transfer or cell-depletion experiments.

These questions define the next stage of the work.

## Conclusions

The work reported here establishes CagW as a workable mRNA vaccine antigen. CagW-Fc mRNA-LNPs raised mucosal and systemic humoral responses, primed Th1/Th2/Th17-type cellular immunity and protected C57BL/6 mice against *H. pylori* challenge while limiting tissue damage; the LNP shell protected the transcript from degradation and improved antigen expression. These data justify further development of mRNA-LNP vaccines for *H. pylori* and other gastrointestinal pathogens, including optimisation of the delivery route and combination of CagW with additional conserved antigens.

## List of abbreviations

ANOVA: analysis of variance
APC: antigen-presenting cell
cagPAI: cag pathogenicity island
CagW: component of the H. pylori type IV secretion system
CFU: colony-forming unit
CTL: cytotoxic T lymphocyte
DC: dendritic cell
ELISA: enzyme-linked immunosorbent assay
ExPASy: Expert Protein Analysis System
Fc: crystallisable fragment of immunoglobulin
GRAVY: grand average of hydropathicity
HE: haematoxylin and eosin
HEK293T: human embryonic kidney 293T cells
IFA: indirect immunofluorescence assay
IFN-γ: interferon-gamma
Ig: immunoglobulin (IgG, IgM, IgA, sIgA, IgG1, IgG2a)
IL: interleukin
LNP: lipid nanoparticle
m1Ψ: N1-methylpseudouridine
MALT: mucosa-associated lymphoid tissue
mRNA: messenger RNA
PDI: polydispersity index
PPI: proton pump inhibitor
PSIPRED: Position-Specific Iterative protein secondary-structure prediction
SP: signal peptide
T4SS: type IV secretion system
TLR: Toll-like receptor
UniProt: Universal Protein Resource

## Declarations

### Authors’ contributions

Yunwang Zhao: original draft preparation; Hongxian Cheng: writing—review and editing; Shiyi Zhou: visualisation; Lei He and Chenlin Xiu: methodology and formal analysis; Baihai Huang: conceptualisation; Xiongxiong Lu: writing-review and editing, original draft preparation, methodology and conceptualisation. All authors read and approved the final manuscript.

## Acknowledgements

The authors gratefully acknowledge financial support from the National Natural Science Foundation of China.

## Availability of data and materials

The datasets generated and analysed during the current study are available from the corresponding author on reasonable request.

## Consent for publication

Not applicable.

## Ethics approval and consent to participate

All animal procedures complied with institutional regulations for the care and use of laboratory animals. The protocol was approved by the Animal Care and Use Committee of Ruijin Hospital, Shanghai Jiao Tong University School of Medicine (approval code 2024Z010) and performed in accordance with its guidelines.

